# Top dressed biochar increases tree seedling growth and decreases sodium leaching

**DOI:** 10.1101/2023.08.03.551785

**Authors:** Brian Wagner, Allyson Salisbury, Meghan G. Midgley

**Affiliations:** Center for Tree Science, The Morton Arboretum, 4100 Illinois Route 53, Lisle, Illinois 60532 USA; Illinois Math and Science Academy, 1500 Sullivan Rd, Aurora, Illinois 60506 USA; University of Notre Dame, Notre Dame, Indiana 46556 USA

**Author notes:** Corresponding author E-mail address, Postal address: Illinois Route 53, Lisle, IL 60532.

**Keywords:** Ecophysiology, mitigation, sodic, soil amendments, urban soil

## Abstract

De-icing salts on roadways are nearly ubiquitous in northern cities during winter months, leading to contamination of soils adjacent to roadways. Sodium chloride salts often have detrimental impacts on water and trees, though some species are more sensitive than others. Biochar has the potential to mitigate sodium’s harmful effects due to its large surface area:volume ratio and subsequent ability to sorb ions from solution. We conducted a four-month greenhouse experiment to test if biochar applied as either a top dressing or incorporated into the growing medium reduced sodium leaching and buffered tree responses to sodium stress. We also evaluated the effects of salt addition and biochar on four tree species that vary in salt tolerance: *Catalpa speciosa* (tolerant), *Gleditsia triacanthos* (tolerant), *Acer saccharum* (intolerant), and *Quercus rubra* (intolerant). We found no interactive effects of sodium addition and biochar on sodium leaching or tree growth and physiology. However, we did find that top dressed biochar broadly decreased sodium leaching, likely via positive effects of top dressed biochar on tree seedling growth, *Catalpa speciosa* in particular. Incorporated biochar, on the other hand, had positive or neutral effects on sodium leaching and negative effects on the production of new shoots and fine roots. Given that biochar is a relatively expensive amendment, it should be used sparingly to improve urban tree growth and health. Overall, this study shows that biochar application decisions have implications for tree growth and soil management.

## INTRODUCTION

Sodium chloride is the most common deicing agent applied to roads in northern regions where ice on roads can pose a danger to drivers (Buttle and Labadia, 1999; Silver et al., 2009; Thunqvist, 2004). Soil contamination via salty runoff from these roads negatively affects the survival, growth, and health of roadside trees (Chaves et al., 2009; Goodrich and Jacobi, 2012; Munns, 2002; Thomas et al., 2013). Salt stress may be particularly acute in urban soils as salt can exacerbate water limitation (Benson et al., 2019; Pitt et al., 2008). Selecting salt-tolerant tree species for roadside plantings may mitigate the negative impacts of sodium chloride on trees (Li et al., 2017; Munns and Tester, 2008; Wu, 2018). However, narrowing species selection to only salt-tolerant species is incompatible with the goal of increasing urban canopy diversity (Alvey, 2006; Morgenroth et al., 2016). Furthermore, sodium ions in soils can also leach out to waterways, thereby extending the effects of salt contamination far downstream of the source (Karraker et al., 2008; Kaushal et al., 2005; Sanzo and Hecnar, 2006). Increasing the diversity and health of roadside trees and decreasing downstream impacts requires decreasing salt availability via soil remediation.

Biochar shows great promise as a soil amendment that can mitigate the negative effects of salt on trees (Di Lonardo et al., 2017; Drake et al., 2016; Somerville et al., 2020; Thomas et al., 2013) and decrease sodium and nutrient leaching from soils (Di Lonardo et al., 2017; Li et al., 2018; Seguin et al., 2018). Biochar increases soil contaminant/toxin sorption, water-holding capacity, and pH, all of which benefit trees (Thomas and Gale, 2015; Yuan et al., 2019). However, the few studies examining biochar’s ability to offset salt effects on trees show mixed results (Di Lonardo et al., 2017; Scharenbroch et al., 2022). In a six month greenhouse experiment, biochar decreased salinity damage to a salt-sensitive tree species, but had no effects on a salt-tolerant tree species (Di Lonardo et al., 2017). Furthermore, in 5-15 year old street trees, biochar addition neither reduced soil sodium concentrations nor enhanced tree growth or chlorophyll contents (Scharenbroch et al., 2022). Biochar may also mitigate water stress induced by road salts. In a field study where biochar was incorporated into urban soils, biochar addition increased plant-available water and tree growth and decreased water stress, but only in sandy soils (Somerville et al., 2020). Thus, despite biochar’s promise for offsetting the negative effects of road salt, current evidence is scant.

In addition to improving tree growth and health by offsetting the negative effects of salt, biochar may also positively impact trees via other mechanisms. While the majority of studies have found an improvement of tree health and growth (Schaffert et al., 2022; Thomas and Gale, 2015), there are differing conclusions about the influence of biochar compared to other organic matter and fertilizer amendments, with some showing that biochar leads to greater growth than other amendments (Scharenbroch et al., 2013), others showing no difference between amendment types (Somerville et al., 2019), and others showing that biochar leads to less growth than other amendments (Yousaf et al., 2021). Furthermore, some studies indicate that biochar’s effects are either enhanced or only effective when applied jointly with fertilizer (Schmidt et al., 2017), compost (Ghosh et al., 2015), or mulch (Percival et al., 2023). Other factors such as plant species and time may also influence the magnitude of biochar’s effects (Fite et al., 2019; Noyce et al., 2017). Finally, biochar effects vary across tissues; in a three month greenhouse study, biochar incorporation into urban soils increased the biomass, height, and chlorophyll contents of silver maple seedlings, but had little effect on root allocation, caliper, or photosynthesis measurements (Sifton et al., 2022). As a result, the effects of biochar itself on tree health and growth are yet to be determined with certainty.

Although much research has been conducted on biochar and its effects on plants and soils, there is a dearth of studies comparing different application strategies. Previous research has focused primarily on tree seedling responses to top dressed biochar (Scharenbroch et al., 2013; Thomas et al., 2013) or biochar incorporation into growing medium (Drake et al., 2016; Li et al., 2022; Piao et al., 2023). Although most studies evaluate just one application strategy, some studies comparing biochar application strategies found no effects on nitrate and ammonium leaching (Li et al., 2018), but variability in root biomass (Zou et al., 2023). Biochar addition broadly increased root biomass relative to unamended soils, but burying a 1 cm-thick horizontal band of biochar into the soil led to a greater increase in biomass than fully incorporating biochar into soil or burying numerous “strips” of biochar under the soil surface at different depths and angles (Zou et al., 2023). As such, biochar effects on trees and its ability to offset salt effects on trees and soils may vary with application strategy.

In this study, we addressed two overarching questions: (1) To what extent does biochar reduce sodium leaching, and (2) does biochar buffer tree responses to sodium addition? We hypothesized that sodium addition would increase sodium leaching and decrease chlorophyll content, photosynthesis, and growth of trees, particularly those that are generally considered to be salt-intolerant. We also hypothesized that biochar addition would generally increase tree chlorophyll content, photosynthesis, and growth. We further hypothesized that biochar addition to soils would offset any effects of sodium addition on leachate chemistry and tree responses.

Specifically, we hypothesized that top dressed biochar would best mediate the effects of added sodium by intercepting salt before it entered the soil system and tree roots. Finally, we hypothesized that salt-intolerant trees would be most negatively impacted by salt addition and thereby benefit most from biochar amendments.

## MATERIAL AND METHODS

We conducted a four month greenhouse experiment to evaluate the effects of biochar amendments and salt addition on sodium leaching and tree seedling physiology and growth. To assess biochar effects, we created three growing mediums: a control soil containing no biochar, a medium where biochar was incorporated into the soil, and a medium where biochar was top dressed. These mediums were added to 3.8L pots (15.2cm diameter, 17.8cm depth) that were externally lined with landscape fabric to prevent medium loss. For the controls, we created a growing medium consisting of v/v 50% sand and 50% commercial garden soil (Nature’s Care Organic Garden Soil, Miracle-Gro, Marysville, OH, USA). For both biochar treatments, biochar represented 5% of the medium volume as recommended by Schaffert et al. (2022) and used by Fite et al. (2019). Biochar was .55mm - 2.0mm in size and derived from Southern Yellow Pine produced by fast pyrolysis at temperatures ranging from 550°C - 900°C (NAKED Char, American BioChar Company, Niles, MI, USA). When biochar was incorporated into the soil, we created a mix of v/v 47.5% sand, 47.5% commercial garden soil, and 5% biochar. When biochar was top dressed, we added the v/v 50% sand and 50% commercial garden soil mix to each pot, but 5% less (by weight) than the controls. We then added 5% biochar to the medium surface (∼1 cm depth).

To assess whether biochar and salt effects varied across species, we planted one of four tree species into each pot: *Catalpa speciosa*, *Gleditsia triacanthos*, *Acer saccharum*, or *Quercus rubra*. These four species were chosen because they vary in salt tolerance and due to their popularity in urban landscaping; *Catalpa speciosa* and *Gleditsia triacanthos* are considered to be salt-resistant species while *Acer saccharum* and *Quercus rubra* are salt-intolerant (https://mortonarb.org/plant-and-protect/search-trees-and-plants/). We purchased 30 to 60 cm tall bare root seedlings of each species (Cold Stream Farm LLC, Free Soil, MI, USA). Before planting, we measured seedling biomass, stem diameter, stem water displacement, and coarse root water displacement. We did not measure stem water displacement for *Gleditsia triacanthos* because the stems were longer than the displacement equipment. Two perpendicular stem diameter measurements were taken at 2.5cm above the root flare; we marked this location with masking tape to ensure initial and final measurements were collected from the same location.

To account for variability in initial tree sizes and greenhouse conditions, we incorporated a block design into our experiment. Our experiment consisted of seven blocks, and each block contained six individuals of each species with two individuals of each species planted into each of the three growing medium (24 pots per block; 168 pots total). Because seedlings of a given species varied in initial sizes, we sorted individuals from each species by height and individuals were split into seven groups containing six similarly-sized individuals of each species. These groups of six were each assigned to a block with the six smallest seedlings of all species assigned to block 1 and the largest six assigned to block 7. Species were clustered within a block. Species locations within each block and growing medium locations within each species cluster were randomly assigned. We established the experiment in mid-May 2019 and allowed seedlings to acclimate and leaf out for four weeks.

On June 17, 2019, we placed each pot into a 17.8cm diameter, 17.8cm depth kids sand bucket that allowed us to collect leachate weekly, and we began adding sodium chloride at a rate of 4g Na^+^ m^-2^. We added sodium chloride solutions to half of the pots in each cluster, one of each growing medium. As such, we added 0.009435g Na^+^ to each “salt addition” pot weekly over the course of eight weeks, approximating salt deposition measurements taken by Kelsey and Hootman (1991) adjacent to the Illinois Tollway. When we initiated the salt addition component of this experiment, we also began watering the trees with uniform volumes of water: 750mL per week. Our weekly salt and water addition and leachate collection protocol was as follows: every Monday, we added 250mL of water to each “no salt” pot and 200mL of water and 50mL of sodium chloride solution to each “salt addition” pot. On Wednesdays, all trees were watered with 250mL of water. Leachate volumes were measured and leachate subsamples were collected from blocks 1-4. This method of sample collection and watering was repeated on Fridays, but leachates were collected from blocks 5-7. Leachate accumulated in the bottom of the pails on days in which it was not collected. We concluded this component of the experiment on August 9, 2019. To measure leachate sodium concentrations, we mixed leachate subsamples with 1% La_2_O_3_ and 0.1% CsCl releasing agents in a 30:1:1 ratio and measured sodium concentrations in these solutions on an atomic adsorption spectrophotometer (AAnalyst 400, Perkin Elmer, Waltham, MA, USA). We calculated the total sodium leached from a pot in a given sampling week using these concentrations and total weekly leachate volume measurements.

During and after the salt addition phase, we collected tree physiology measurements. We measured leaf chlorophyll contents at the end of week six of the salt addition phase and three weeks after the conclusion of salt addition using a SPAD meter (SPAD 502 Plus, Konica Minolta Inc., Tokyo, Japan). When available, we collected SPAD measurements on three leaves from each of the *Catalpa speciosa*, *Acer saccharum*, and *Quercus rubra* trees and 5 leaves from *Gleditsia triacanthos* trees and calculated mean SPAD. When fewer leaves were available, we collected data from all available leaves. We averaged these two temporal SPAD measurements for statistical analyses. Between the beginning of week seven and week eight of the salt addition phase, we measured photosynthesis rates (LI-6800 Portable Photosynthesis System, LI-COR, Lincoln, NE, USA). We collected measurements from all trees of a given species on the same day. Because *Gleditsia triacanthos* leaves and one *Quercus rubra* leaf did not fill the area of the chamber surface, we photographed the measured leaves and calculated leaf area using Image J (Schneider et al., 2012). Due to a combination of leaf browning and dieback, we collected week six SPAD measurements on 126 of the original 168 seedlings, final SPAD measurements on 122 seedlings, and photosynthesis measurements on 127 seedlings.

At the conclusion of the experiment in mid-September 2019, we destructively harvested the seedlings and took a variety of measurements to assess salt and biochar effects on tree seedling growth. To quantify changes in the growth of pre-existing tissues, we re-measured stem diameter and stem and coarse root water displacement as above. We calculated growth based on water displacement and biomass of pre-existing tissues and diameter using initial and final measurements. To quantify the changes in biomass of pre-existing tissues and biomass of new tissues, we clipped newly produced shoots and leaves off of the pre-existing stems and newly produced fine roots (<2mm diameter) off the pre-existing coarse roots. We separately dried the stems, coarse roots, new shoots, and fine roots at 55°C for one week before weighing them. We calculated changes in preexisting tissue and total biomass using final and initial weight data.

We used a series of mixed linear models to evaluate the effects of biochar amendments and salt addition on sodium leaching and tree seedling physiology, growth, and changes in biomass. To assess temporal changes in sodium leaching over the course of the salt addition phase, we created a model with total sodium leached as a function of week as a fixed effect and block as a random effect. We evaluated the effects of biochar amendments (control, biochar incorporated, or biochar topdressing), sodium addition (no salt or salt addition), their interaction, and species on weekly sodium leaching, average SPAD, photosynthesis rates, and growth and biomass metrics using block as a random effect. Finally, we subset the data by species to examine species-specific SPAD, photosynthesis, and growth and biomass responses to biochar amendments, salt, and their interaction, maintaining block as a random effect. We transformed the data as necessary to meet residual normality and heteroskedasticity assumptions. Results were considered significant if they had *p*-values < 0.05. When we found a significant main effect of amendments, salt by amendment interaction, or species, we conducted a Tukey HSD post-hoc test to identify differences among groups. Model statistics are reported in tables and post-hoc *p*-values are reported in the text. All statistical analyses were conducted in R using the lmerTest package for mixed linear models (Kuznetsova et al. 2017) and lsmeans package for post-hoc tests (Lenth 2016). Figures were created using ggplot2 (Wickham 2016).

## RESULTS

### Sodium leaching

Sodium leaching decreased over the course of the 8 week experiment and varied among weeks (F_(7,1315)_ = 332.42, *p* < 0.001 ; Figure 1). More sodium leached in week 1 than in any subsequent week (*p* < 0.001). Sodium leached during the second week of the experiment was also significantly greater than that of each following week (*p* < 0.001), and weeks 3, 4, and 6 each had significantly greater sodium leaching than week 8 (*p* ≤ 0.03).

**Figure 1.**
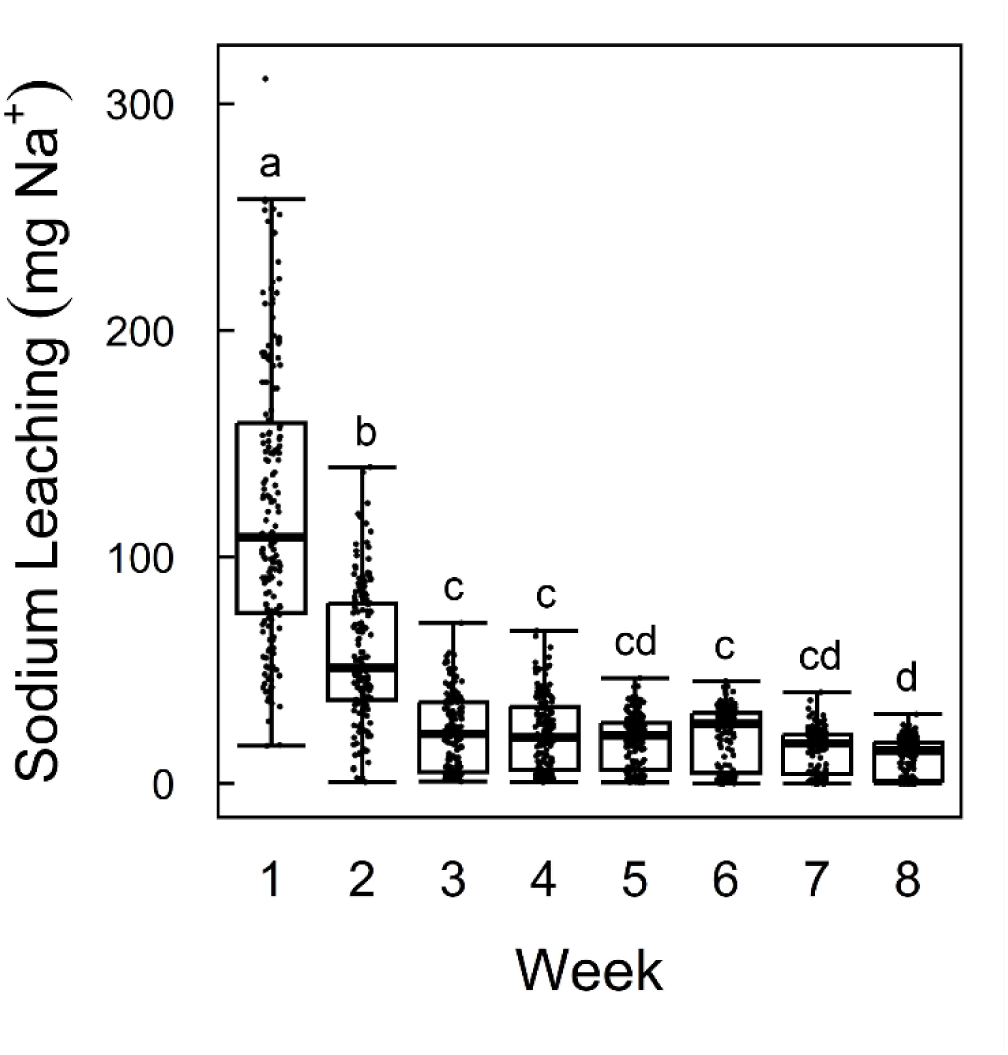
Box and whisker plot showing sodium leaching (in mg) of all trees by week. The thick horizontal line inside each box represent the median value, the box represents the 25-75 percentile, and the whiskers extend from the first quartile minus 1.5*the interquartile range to the third quartile plus 1.5*the interquartile range. Each point on the graph is an individual tree’s sodium leachate measurement from that week. Unique significance letters indicate statistically different mean sodium leachate masses per Tukey’s HSD test (*p* < 0.05).

Sodium leaching varied among soil treatments and tree species and with sodium addition (Table 1; Figure 2). In week 1, we found that more sodium leached from biochar incorporated pots than from controls (*p* = 0.03; Figure 2a). In contrast, in weeks 3, 4, 6, 7, and 8 less sodium leached from biochar top dressed pots than from controls (*p* ≤ 0.03; Figure 2d) while in week 8 less sodium also leached from biochar top dressed pots than from biochar incorporated pots (*p* = 0.01; Figure 2g).

**Figure 2.**
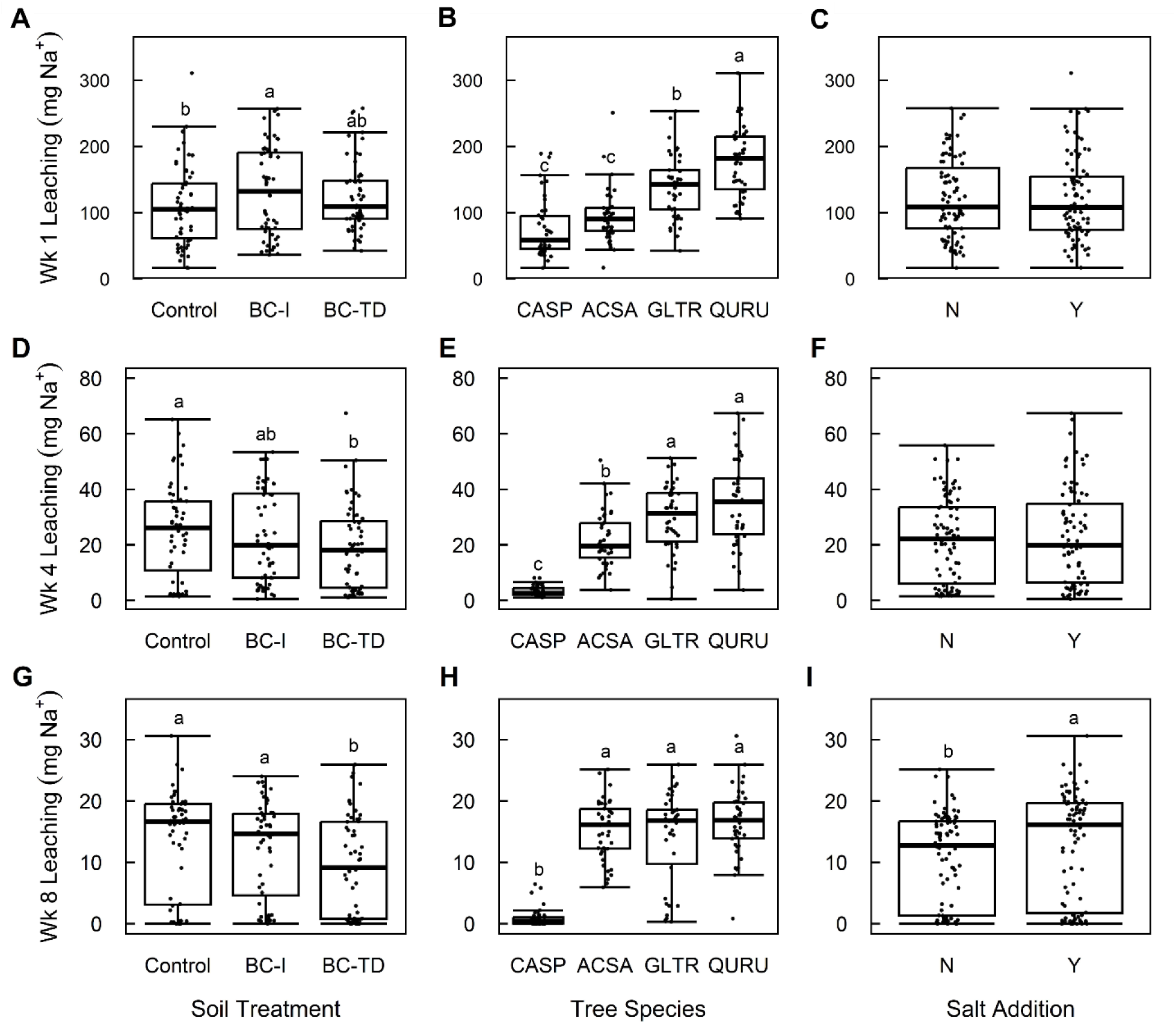
Box and whisker plots of sodium leachate for weeks 1 (**A**, **B**, and **C**), 4 (**D**, **E**, and **F**), and 8 (**G**, **H**, and **I**) by soil treatment (**A**, **D**, and **G**), tree species (**B**, **E**, and **G**), and salt addition (**C**, **F**, and **I**). The thick horizontal line inside each box represents the median value, the box represents the 25-75 percentile, and the whiskers extend from the first quartile minus 1.5*the interquartile range to the third quartile plus 1.5*the interquartile range. Each point on the graph is an individual tree’s sodium leachate measurement from that week. Control = control soils (no biochar), BC-I = incorporated biochar, BC-TD = top dressed biochar, CASP = *Catalpa speciosa*, ACSA = *Acer saccharum*, GLTR = *Gleditsia triacanthos*, QURU = *Quercus rubra*, N = no salt treatment, Y = salt treatment.

**Table 1.** F and *p* values for various sodium leachate, photosynthesis, SPAD, and biomass growth responses to soil treatment (control, top dressed biochar, and incorporated biochar), tree species (*Acer saccharum*, *Catalpa speciosa*, *Gleditsia triacanthos*, and *Quercus rubra*), salt treatment (no salt added and salt added), and interaction of soil and salt treatments.

| Response | Soil treatments | Tree species | Salt addition | Soil x salt |
| --- | --- | --- | --- | --- |
| Week 1 sodium leaching (mg Na<sup>+</sup>) | F<sub>2,158</sub> = 3.24,<br>$p = 0.04$ | F<sub>3,158</sub> = 39.78,<br>$p < 0.001$ | F<sub>1,158</sub> = 0.01,<br>$p = 0.94$ | F<sub>2,158</sub> = 0.66,<br>$p = 0.52$ |
| Week 2 sodium leaching (mg Na<sup>+</sup>) | F<sub>2,148.09</sub> = 2.52,<br>$p = 0.08$ | F<sub>3,148.09</sub> = 72.54,<br>$p < 0.001$ | F<sub>1,148.06</sub> = 0.08,<br>$p = 0.78$ | F<sub>2,148.11</sub> = 1.53,<br>$p = 0.22$ |
| Week 3 sodium leaching (mg Na<sup>+</sup>) | F<sub>2,139.06</sub> = 3.40,<br>$p = 0.04$ | F<sub>3,139.54</sub> = 71.88,<br>$p < 0.001$ | F<sub>1,139.09</sub> = 0.05,<br>$p = 0.82$ | F<sub>2,139.22</sub> = 0.71,<br>$p = 0.50$ |
| Week 4 sodium leaching (mg Na<sup>+</sup>) | F<sub>2,153</sub> = 4.55,<br>$p = 0.01$ | F<sub>3,153</sub> = 70.74,<br>$p < 0.001$ | F<sub>1,153</sub> = 0.41,<br>$p = 0.52$ | F<sub>2,153</sub> = 1.24,<br>$p = 0.29$ |
| Week 5 sodium leaching (mg Na<sup>+</sup>) | F<sub>2,153</sub> = 2.54,<br>$p = 0.08$ | F<sub>3,153</sub> = 112.39,<br>$p < 0.001$ | F<sub>1,153</sub> = 1.58,<br>$p = 0.21$ | F<sub>2,153</sub> = 0.11,<br>$p = 0.90$ |
| Week 6 sodium leaching (mg Na<sup>+</sup>) | F<sub>2,158</sub> = 4.36,<br>*p* = 0.01 | F<sub>3,158</sub> = 119.53,<br>*p* < 0.001 | F<sub>1,158</sub> = 0.26,<br>*p* = 0.61 | F<sub>2,158</sub> = 0.42,<br>*p* = 0.66 |
| Week 7 sodium leaching (mg Na<sup>+</sup>) | F<sub>2,159</sub> = 5.22,<br>*p* = 0.01 | F<sub>3,159</sub> = 97.93,<br>*p* < 0.001 | F<sub>1,159</sub> = 2.25,<br>*p* = 0.14 | F<sub>2,159</sub> = 2.35,<br>*p* = 0.10 |
| Week 8 sodium leaching (mg Na<sup>+</sup>) | F<sub>2,153</sub> = 9.20,<br>*p* < 0.001 | F<sub>3,153</sub> = 94.46,<br>*p* < 0.001 | F<sub>1,153</sub> = 7.62,<br>*p* = 0.01 | F<sub>2,153</sub> = 0.93,<br>*p* = 0.40 |
| SPAD | F<sub>2,119.86</sub> = 0.46,<br>*p* = 0.63 | F<sub>3,119.76</sub> = 42.46,<br>*p* < 0.001 | F<sub>1,119.62</sub> = 0.26,<br>*p* = 0.61 | F<sub>2,119.77</sub> = 0.56,<br>*p* = 0.57 |
| Photosynthesis rate (μmol C m<sup>-2</sup> s<sup>-1</sup>) | F<sub>2,111.97</sub> = 2.62,<br>*p* = 0.08 | F<sub>3,112.61</sub> = 42.13,<br>*p* < 0.001 | F<sub>1,111.78</sub> = 0.21,<br>*p* = 0.65 | F<sub>2,111.01</sub> = 0.56,<br>*p* = 0.57 |
| Diameter growth (%) | F<sub>2,158</sub> = 1.58,<br>*p* = 0.21 | F<sub>3,158</sub> = 28.18,<br>*p* < 0.001 | F<sub>1,158</sub> = 1.00,<br>*p* = 0.32 | F<sub>2,158</sub> = 1.57,<br>*p* = 0.21 |
| Stem growth (%) | F<sub>2,117</sub> = 1.60,<br>*p* = 0.21 | F<sub>2,117</sub> = 68.55,<br>*p* < 0.001 | F<sub>1,117</sub> = 0.90,<br>*p* = 0.34 | F<sub>2,117</sub> = 0.46,<br>*p* = 0.64 |
| Coarse root growth (%) | F<sub>2,151.96</sub> = 5.06,<br>*p* = 0.01 | F<sub>3,151.96</sub> = 71.40,<br>*p* < 0.001 | F<sub>1,151.96</sub> = 0.63,<br>*p* = 0.43 | F<sub>2,151.96</sub> = 2.81,<br>*p* = 0.06 |
| New shoot biomass (g) | F<sub>2,153</sub> = 3.88,<br>*p* = 0.02 | F<sub>3,153</sub> = 61.18,<br>*p* < 0.001 | F<sub>1,153</sub> = 0.73,<br>*p* = 0.40 | F<sub>2,153</sub> = 1.30,<br>*p* = 0.28 |
| New fine root biomass (g) | F<sub>2,152.12</sub> = 4.03,<br>*p* = 0.02 | F<sub>3,152.12</sub> = 30.42,<br>*p* < 0.001 | F<sub>1,152.12</sub> = 2.15,<br>*p* = 0.14 | F<sub>2,152.12</sub> = 1.35,<br>*p* = 0.26 |
| Stem and coarse root biomass (%) | F<sub>2,152</sub> = 7.71,<br>*p* = 0.001 | F<sub>3,152</sub> = 158.95,<br>*p* < 0.001 | F<sub>1,152</sub> = 0.27,<br>*p* = 0.60 | F<sub>2,152</sub> = 1.78,<br>*p* = 0.17 |
| New shoot and fine root biomass (g) | F<sub>2,152.09</sub> = 4.51,<br>*p* = 0.01 | F<sub>3,152.09</sub> = 56.15,<br>*p* < 0.001 | F<sub>1,152.09</sub> = 1.33,<br>*p* = 0.25 | F<sub>2,152.09</sub> = 1.50,<br>*p* = 0.23 |
| Total biomass (%) | F<sub>2,152.01</sub> = 4.92,<br>*p* = 0.01 | F<sub>3,152.01</sub> = 115.19,<br>*p* < 0.001 | F<sub>1,152.01</sub> < 0.01,<br>*p* = 0.94 | F<sub>2,152.01</sub> = 1.10,<br>*p* = 0.34 |

Sodium leaching also varied among tree species. More sodium leached from *Quercus rubra* pots than from *Catalpa speciosa* pots in all weeks of the experiment (*p* < 0.001). Furthermore, *Catalpa speciosa* pots consistently had lower sodium leaching than all other species in weeks 2-8 (*p* < 0.001); in week 1, *Catalpa speciosa* and *Acer saccharum* pots exhibited similar sodium leaching (Figure 2b). *Gleditsia triacanthos* and *Quercus rubra* pots had similarly high sodium leaching aside from weeks 1 and 6 where more sodium leached from *Quercus rubra* pots than *Gleditsia triacanthos* pots (*p* ≤ 0.01; Figure 2b). *Acer saccharum* pots exhibited intermediate sodium leaching during the middle of the experiment with lower sodium leaching than *Quercus rubra* and *Gleditsia triacanthos* pots during weeks 2, 3, 4, and 5 of the experiment (*p* ≤ 0.01; Figure 2e). However, by week 7 and 8 of the experiment, *Acer saccharum, Quercus rubra*, and *Gleditsia triacanthos* pots leached similarly high amounts of sodium (Figure 2h).

Sodium leaching was also greater in sodium addition pots, but only towards the end of the experiment. In weeks 1-7, we found no effects of sodium addition on sodium leaching (Figure 2c,f). We also found no interactions between sodium addition and soil amendments. However, in week 8, more sodium leached from sodium addition pots than from control pots (Figure 2i).

### SPAD and Photosynthesis

SPAD and photosynthesis rates varied among species, but treatment effects were species-specific (Table 1; Figure 3). *Acer saccharum*, *Gleditsia triacanthos*, and *Quercus rubra* had higher SPAD than *Catalpa speciosa* (*p* < 0.001; Figure 3a). However, *Catalpa speciosa* was the only species that responded to soil treatments (Table 2); *Catalpa speciosa* growing in biochar top dressed pots had higher SPAD than *Catalpa speciosa* growing in control or biochar incorporated pots (*p* < 0.001; Figure 3b). *Catalpa speciosa*, *Gleditsia triacanthos*, and *Quercus rubra* had higher photosynthesis rates than *Acer saccharum* (*p* < 0.001; Figure 3c). In this case, only *Acer saccharum* responded to treatments (Supplementary Table 1); *Acer saccharum* photosynthesis rates exhibited an interaction between sodium addition and soil amendments (Figure 3d). *Acer saccharum* photosynthesis rates tended to be greatest when grown in biochar top dressed pots compared to individuals grown in biochar incorporated pots, but only when no salt was added (*p* = 0.11).

**Figure 3.**
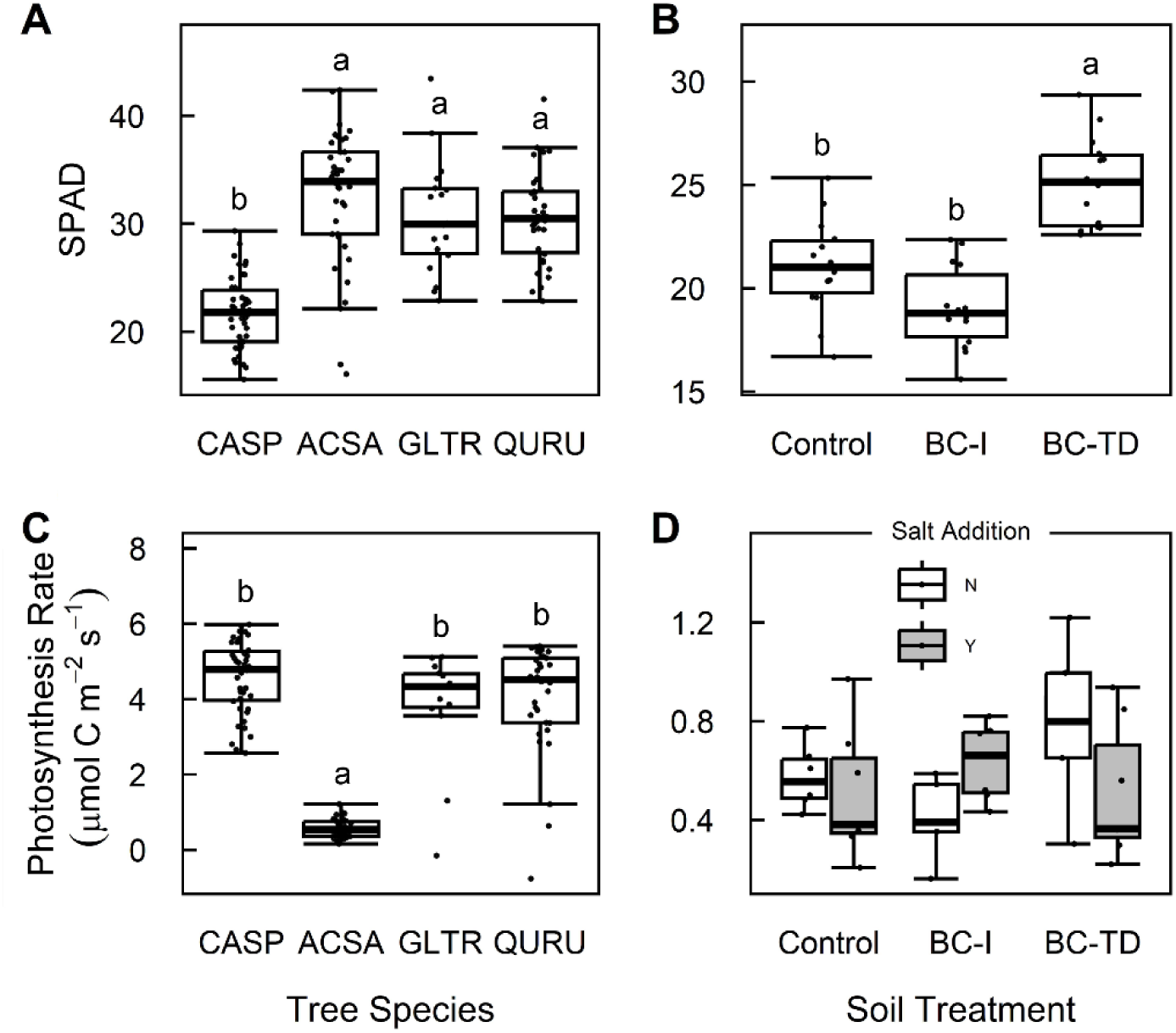
Box and whisker plots of SPAD (**A** and **B**) and photosynthesis rate (**C** and **D**) by tree species (**A** and **C**) and soil treatment (**B** and **D**). The thick horizontal line inside each box represents the median value, the box represents the 25-75 percentile, and the whiskers extend from the first quartile minus 1.5*the interquartile range to the third quartile plus 1.5*the interquartile range. Each point on the graph is an individual tree’s SPAD or photosynthesis rate value. Control = control soils (no biochar), BC-I = incorporated biochar, BC-TD = top dressed biochar, CASP = *Catalpa speciosa*, ACSA = *Acer saccharum*, GLTR = *Gleditsia triacanthos*, QURU = *Quercus rubra*.

**Table 2.** F and *p* values for soil treatment (control, top dressed biochar, and incorporated biochar), salt treatment (no salt added and salt added), and interaction of soil and salt treatments for a variety of *Catalpa speciosa* measurements.

| Response | Soil treatments | Salt addition | Soil x salt |
| --- | --- | --- | --- |
| SPAD | $F_{2,36} = 27.05$,<br>$p < 0.001$ | $F_{1,36} = 0.82$,<br>$p = 0.37$ | $F_{2,36} = 0.03$,<br>$p = 0.97$ |
| Photosynthesis rate<br>($\mu\text{mol C m}^{-2} \text{ s}^{-1}$) | $F_{2,30} = 1.97$,<br>$p = 0.16$ | $F_{1,30} = 0.19$,<br>$p = 0.66$ | $F_{1,30} = 0.85$,<br>$p = 0.44$ |
| Diameter growth (%) | $F_{2,30} = 1.03$,<br>$p = 0.37$ | $F_{1,30} = 0.01$,<br>$p = 0.93$ | $F_{2,30} = 0.63$,<br>$p = 0.54$ |
| Stem growth (%) | $F_{2,36} = 4.00$<br>$p = 0.03$ | $F_{1,36} = 0.47$,<br>$p = 0.50$ | $F_{2,36} = 0.89$,<br>$p = 0.42$ |
| Coarse root growth (%) | $F_{2,30} = 4.41$,<br>$p = 0.02$ | $F_{1,30} = 1.68$,<br>$p = 0.20$ | $F_{2,30} = 1.85$,<br>$p = 0.18$ |
| New shoot biomass (g) | $F_{2,30} = 4.57$,<br>$p = 0.02$ | $F_{1,30} = 0.02$,<br>$p = 0.89$ | $F_{2,30} = 2.70$,<br>$p = 0.08$ |
| New fine root biomass<br>(g) | $F_{2,29.20} = 6.42$,<br>$p = 0.005$ | $F_{1,29.20} = 0.03$,<br>$p = 0.86$ | $F_{2,29.20} = 2.83$,<br>$p = 0.08$ |
| Stem and coarse root<br>biomass (%) | $F_{2,30} = 8.20$,<br>$p = 0.001$ | $F_{1,30} = 0.14$,<br>$p = 0.71$ | $F_{2,30} = 1.28$,<br>$p = 0.29$ |
| New shoot and fine root<br>biomass (g) | $F_{2,29.17} = 5.80$,<br>$p = 0.01$ | $F_{1,29.18} < 0.001$,<br>$p = 1.00$ | $F_{2,29.17} = 3.05$,<br>$p = 0.06$ |
| Total biomass (%) | $F_{2,30} = 5.21$,<br>$p = 0.01$ | $F_{1,30} = 0.09$,<br>$p = 0.76$ | $F_{2,30} = 1.32$,<br>$p = 0.28$ |

### Growth and Changes in Biomass

Growth varied among species and was affected by biochar amendments, but was not impacted by salt addition (Table 1; Figure 4). There was a significant species effect on changes in total biomass, with *Catalpa speciosa* exhibiting the greatest increase in biomass and *Gleditsia triacanthos*, *Quercus rubra*, and *Acer saccharum* having similarly low changes in biomass (*p* < 0.001). Furthermore, changes in tree seedling biomass were greater when biochar was applied as a top dressing than when no biochar was added or biochar was incorporated into the growing medium (*p* ≤ 0.03). This was largely driven by soil treatment effects on *Catalpa speciosa* (Table 2; Figure 5). However, neither salt addition nor the interaction of salt addition and soil type had a significant effect on any growth or biomass measurements (Figures 4c, 4f).

**Figure 4.**
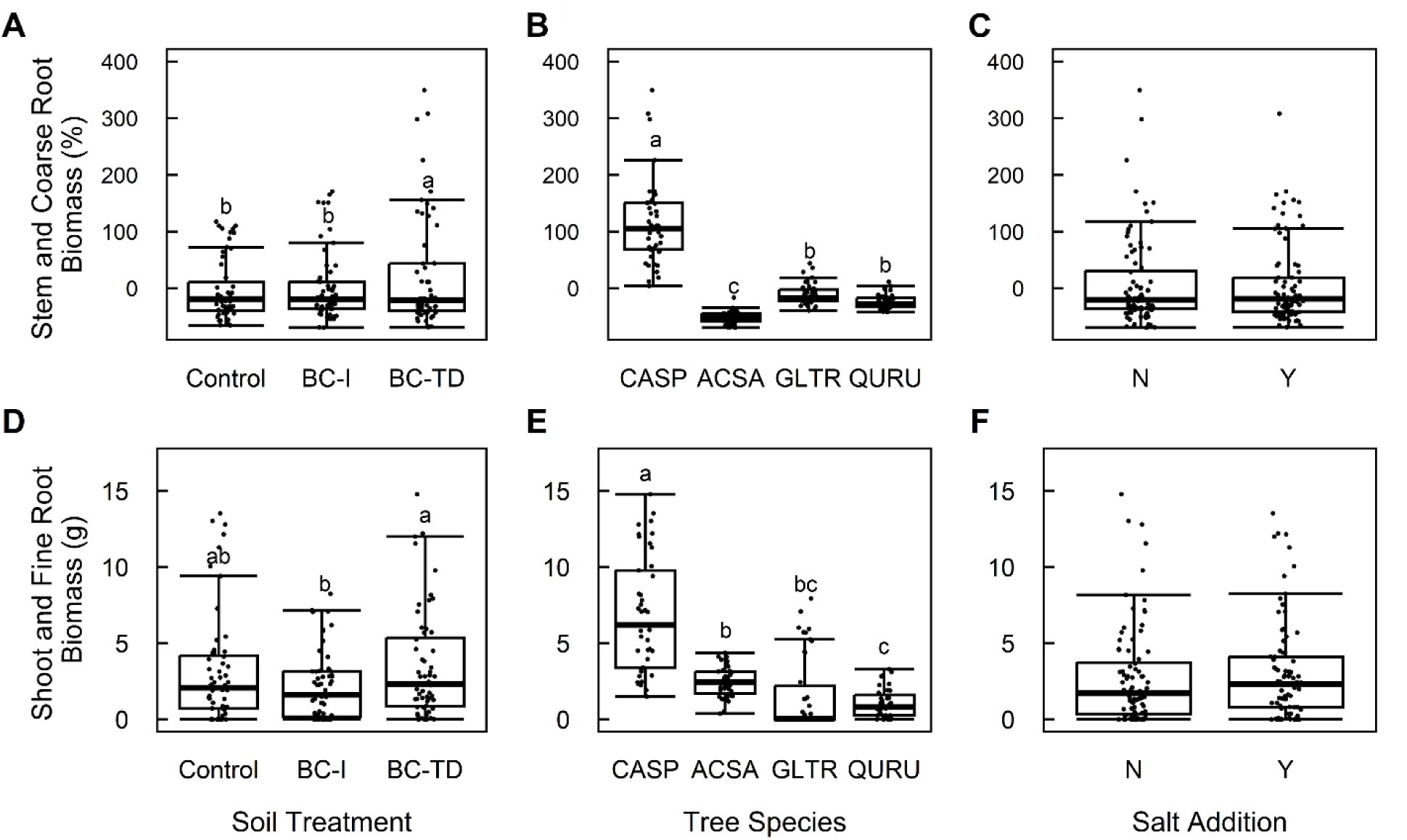
Box and whisker plots of the percent changes in stem and coarse root biomass (**A** - **C**) and the shoot and fine root biomass (**D** - **F**) by soil treatment (**A** and **D**), tree species (**B** and **E**), and salt addition (**C** and **F**). The thick horizontal line inside each box represents the median value, the box represents the 25-75 percentile, and the whiskers extend from the first quartile minus 1.5*the interquartile range to the third quartile plus 1.5*the interquartile range. Each point on the graph is an individual tree’s value. Control = control soils (no biochar), BC-I = incorporated biochar, BC-TD = top dressed biochar, CASP = *Catalpa speciosa*, ACSA = *Acer saccharum*, GLTR = *Gleditsia triacanthos*, QURU = *Quercus rubra*, N = no salt treatment, Y = salt treatment.

**Figure 5.**
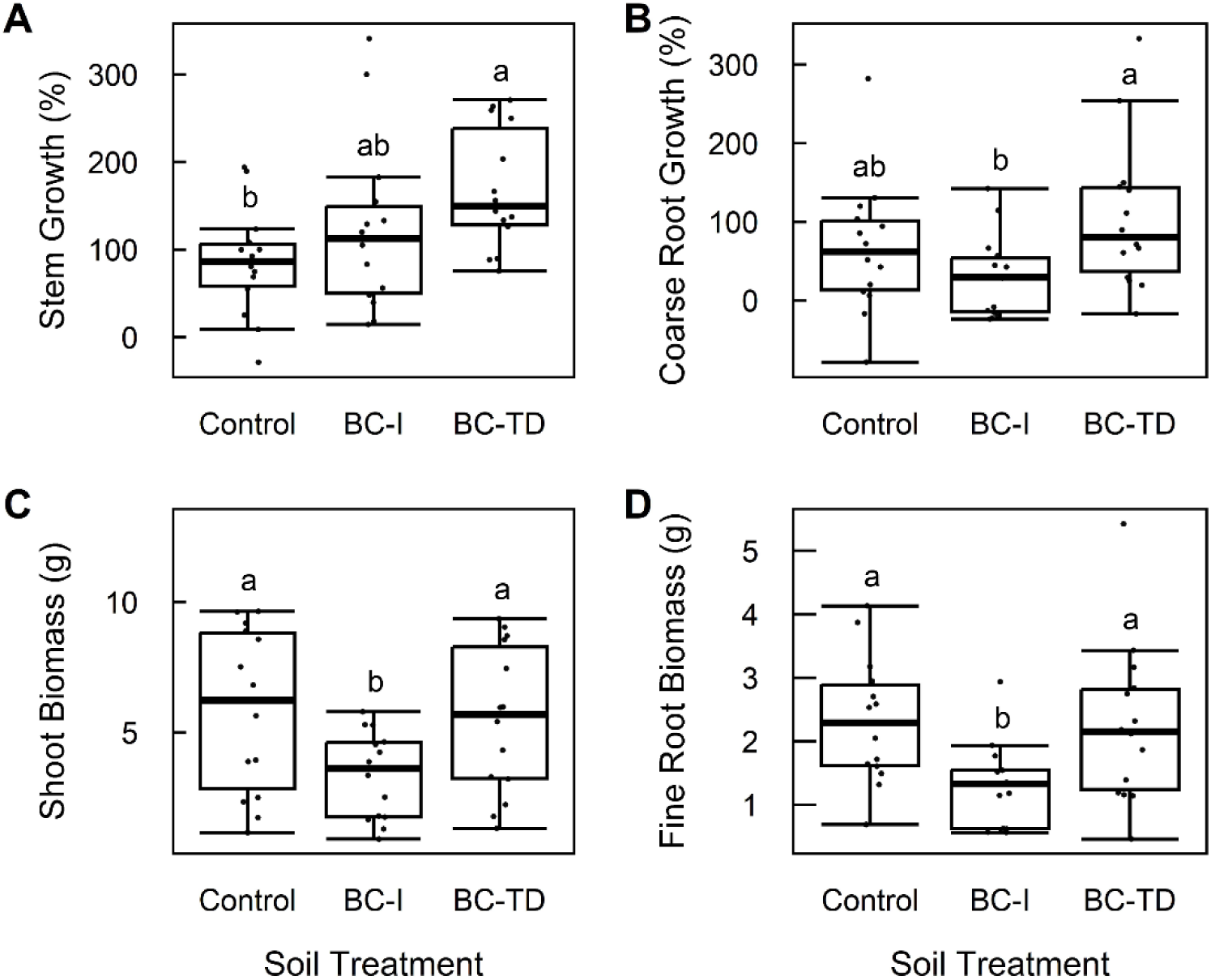
Box and whisker plots of *Catalpa speciosa*’s percent stem growth (**A**), percent coarse root growth (**B**), shoot biomass (**C**), and fine root biomass (**D**) by soil treatment. The thick horizontal line inside each box represents the median value, the box represents the 25-75 percentile, and the whiskers extend from the first quartile minus 1.5*the interquartile range to the third quartile plus 1.5*the interquartile range. Each point on the graph is an individual tree’s sodium leachate measurement from that week. Control = control soils (no biochar), BC-I = incorporated biochar, BC-TD = top dressed biochar.

*Catalpa speciosa* had greater growth and increases in biomass than all other species according to all metrics (*p* < 0.001; Figure 4b; Figure 4e). With regards to pre-existing tissues, *Gleditsia triacanthos* and *Quercus rubra*, had greater increases in biomass than *Acer saccharum* (*p* ≤ 0.01; Figure 4b). Similarly, *Gleditsia triacanthos* coarse root growth was greater than both *Quercus rubra* and *Acer saccharum* coarse root growth, and *Quercus rubra* coarse root growth was greater than that of *Acer saccharum* (*p* ≤ 0.03). In contrast, *Acer saccharum* had greater increases in diameter than *Quercus rubra* (*p* < 0.05). With regards to new tissues, *Acer saccharum* produced more new tissue than *Quercus rubra* (*p* = 0.03; Figure 4e) which was largely driven by greater production of fine roots (*p* = 0.04).

Although soil type did not significantly impact diameter growth or stem growth, it significantly impacted coarse root growth, new above ground biomass, and fine root biomass (Table 1). For all these tissues, top dressed biochar led to greater growth and increases in biomass when compared to biochar incorporated into the growing medium (p ≤ 0.03). Overall, changes in biomass of pre-existing stem and root tissues were greater for top dressed trees than for either biochar incorporated or control trees (*p* ≤ 0.01; Figure 4a) while new tissue biomass was greater when biochar was top dressed than when it was incorporated into the growing medium (*p* = 0.01; Figure 4d).

Tree seedling responses to soil treatments primarily manifested in *Catalpa speciosa* (Table 2; Figure 5). *Catalpa speciosa* responses to top dressed biochar were frequently distinct from *Catalpa speciosa* responses to incorporated biochar while differences between top dressed biochar and controls were less pronounced. Overall, changes in *Catalpa speciosa* biomass were greater when biochar was applied as a top dressing than when biochar was incorporated into the growing medium (*p* = 0.01). In terms of pre-existing tissues, top dressed biochar resulted in greater increases in biomass than incorporated biochar or controls (*p* = 0.005). Top dressed biochar increased stem growth relative to controls (Figure 5a) and increased coarse root growth relative to incorporated biochar (Figure 5b). New tissues responded more consistently to soil treatments with incorporated biochar resulting in lower new shoot biomass, new fine root biomass, and lower new biomass overall compared to top dressed biochar and controls (*p* ≤ 0.05; Figures 5c, 5d).

Other species exhibited limited responses to treatments. *Acer saccharum* trees only showed fine root responses to salt addition, with salt addition leading to greater fine root biomass (Supplementary Table 1; Supplementary Figure 1). Within *Gleditsia triacanthos* trees, the only significant influence was soil type on changes in total biomass (Supplementary Table 2; Supplementary Figure 2), with top dressed trees growing marginally more than those with biochar incorporated into the soil or controls (*p* = 0.07). There were no significant influences of soil type or salt addition on *Quercus rubra* growth or biomass metrics (Supplementary Table 3; Supplementary Figure 3).

## DISCUSSION

Roadside soils can have inherently high levels of sodium (Cunningham et al., 2007; Hootman et al., 1994). Previous studies have shown that incorporating biochar into the growing medium of plants increases leaf chlorophyll concentration and plant growth (Drake et al., 2016; Piao et al., 2023; Ran et al., 2020, p. 20; Sifton et al., 2022; Somerville et al., 2020; Yousaf et al., 2021) and may sorb salts thereby offsetting the harmful effects of sodium on trees and soils (Drake et al., 2016; Thomas et al., 2013). At the outset of this study, we sought to evaluate biochar’s capacity to reduce sodium leaching and buffer tree responses to sodium addition. Due to high sodium concentrations in our growing medium, sodium addition resulted in minimal sodium leaching and had no impacts on tree seedling growth or physiology. As such, biochar did not buffer any negative effects of sodium addition on trees and seemingly had no direct effects on sodium in leachate. However, top dressed biochar decreased sodium leaching and increased tree growth and chlorophyll content, particularly for *Catalpa speciosa.* Rather than buffering the impacts of sodium addition on salt-intolerant trees, biochar addition boosted salt-tolerant tree growth and decreased sodium leaching, potentially by inducing increased tree uptake of sodium. *Sodium addition had minimal effects on sodium leaching or trees*

Surprisingly, sodium chloride addition had minimal effects on sodium leaching and tree growth and physiology. In fact, we detected no salt addition effects on sodium leaching before the final week of sodium chloride addition. This is likely due to the high background levels of sodium in our growing medium that obscured any effects of added salt. Unbeknownst to us at the outset of this experiment, the growing medium we used was inherently sodium-rich, possibly due to some combination of sodium associated with glass in our sand or contamination of the commercial garden soil we used. By week eight, after many weeks of “flushing out” the pre-existing sodium, the salt treatment led to increased sodium leachate compared to the control. However, because salt addition did not have an effect until the 8^th^ and final week of the salt addition phase of the experiment, there was little effect of salt addition on trees. Consequently, we detected no clear interactions between sodium addition and soil treatments.

### Top dressed biochar decreased sodium leaching

Despite minimal effects of sodium addition on sodium leaching, biochar decreased sodium leaching, but only when applied as a top dressing. This is surprising because we would expect biochar to decrease sodium leaching from sodic soils when it is incorporated into the soil thereby maximizing sorption of sodium cations (Drake et al., 2016; Li et al., 2018; Seguin et al., 2018; Thomas and Gale, 2015) although biochar incorporation into surface soils had minimal effects on soil sodium concentrations under 5-15 year old street trees (Scharenbroch et al., 2022). It is unlikely that the top dressed biochar sorbed sodium from the growing medium as there were minimal physical interactions between the two strata. Instead, we suspect that this surprising result is driven by the interaction between top dressed biochar and *Catalpa speciosa* growth. Additionally, it is possible that, compared to control soils, the top dressing of biochar directly influenced the trees themselves to uptake more sodium (Drake et al., 2016), thus preventing that sodium from leaching out of the soil column.

In our experiment, tree species had a much more dramatic effect on sodium leaching than biochar or salt addition. Specifically, *Catalpa speciosa* markedly decreased sodium leaching compared to the other tree species and also put on the most biomass over the course of this experiment, suggesting that *Catalpa speciosa* was directly taking up sodium and storing sodium in its biomass. *Catalpa speciosa* is a salt-tolerant tree, though the particular mechanism of tolerance is unknown. Mechanisms by which other species minimize salt stress of environmentally high concentrations of sodium include prevention of sodium uptake and compartmentalization of cytosolic sodium into vacuoles (Chaves et al., 2009; Wu, 2018). Top dressed biochar increased *Catalpa speciosa* stem and coarse root growth and SPAD relative to incorporated biochar and control treatments, which coincides with the sodium leaching responses. As such, the positive effects of top dressed biochar on *Catalpa speciosa* growth may have indirectly led to a reduction in sodium leaching.

### Biochar application strategy had divergent effects on tree growth

Though positive effects of top dressed biochar on tree seedling growth and changes in biomass were driven by *Catalpa speciosa*, these patterns were generally found across tree species. This finding begs two questions: 1) Why did top dressed biochar increase the growth of pre-existing tissues, and 2) why did incorporated biochar decrease the growth of new tissues?

New shoot and fine root growth of tree seedlings were lower in the biochar incorporated treatment than those of trees in top dressed biochar or control soils. This is likely due to biochar’s ability to sorb nutrients, including plant-available forms of nitrogen such as nitrate from soils (Drake et al., 2016; Shi et al., 2020; Thomas and Gale, 2015). New tissues require a large amounts of nitrogen, so it is possible that tree seedling growth was decreased in the incorporated biochar treatment compared to other soil types due to a biochar-mediated decrease in available nitrate in the root zone, per Liebig’s Law of the Minimum (von Liebig, 1863). This would be in accordance with previous findings in which biochar’s positive effects were enhanced or present only when applied with both organic and inorganic fertilizers to overcome the sorption of pre-existing nutrients (Ghosh et al., 2015; Schmidt et al., 2017). Alternatively, biochar incorporation is correlated with greater sodium uptake by plants when sodium concentration is high (Drake et al., 2016). Therefore, biochar incorporation may have induced sodium uptake in trees and subsequently reduced new biomass production.

On the other hand, our results showed that top dressed biochar increased growth of coarse roots and stems compared to control and biochar incorporated soils, indicating a direct positive effect of top dressed biochar on pre-existing tissue growth in contrast with a negative effect of incorporated biochar on new tissue growth. This contrasts with the findings of (Zou et al., 2023), which found that incorporated biochar increased leaf and root biomass more than localized biochar application strategies did. However, our results are not unique; other greenhouse studies have also found that top dressed biochar increases plant growth (Scharenbroch et al., 2013; Thomas et al., 2013). Because biochar’s direct interaction with the soil is minimized in the top dressed treatments in our experiment, a possible explanation of why the top dressed soil treatments led to increased biomass is that the positive effects of biochar prevail while the negative effects (i.e., sorption of required nutrients) were not expressed. Regardless of mechanism, our experiment clearly demonstrates that biochar application strategies impact tree growth and physiology.

## CONCLUSIONS

Biochar is frequently hailed as a solution to a wide variety of problems including climate change due to its slow decomposition rate and soil infertility as a component of slow-release fertilizer (McBeath et al., 2014; Scharenbroch et al., 2013; Thomas and Gale, 2015; Zou et al., 2023). It has recently been touted as a soil amendment that can increase crop production in saline and sodic soils due to its sorptive properties (Li et al., 2022; Piao et al., 2023; Ran et al., 2020) which may extend to offsetting the negative effects of road salt on roadside trees and water quality (Somerville et al., 2020). Our study demonstrates that biochar effects on sodium leaching from sodic soils or due to sodium addition may be most impacted by indirect effects of biochar on tree growth and physiology rather than direct effects of biochar on sodium sorption. Furthermore, top dressed biochar produced the most desirable outcomes, which is encouraging given that top dressed biochar can be applied quickly and easily to both new and established trees. However, these benefits manifested most in a fast-growing, salt-tolerant tree species, indicating that biochar itself may not be a great solution for decreasing downstream effects of deicing salt on waterways or buffering salt-sensitive tree species responses to road salt inundation. Further studies are needed to assess the ability of biochar to remediate sodic soils and offset road salt effects on trees and waterways in real-world situations (Schaffert et al., 2022). Our study also adds to a growing body of literature demonstrating that the benefits of biochar for trees are context-dependent (Di Lonardo et al., 2017; Fite et al., 2019; Noyce et al., 2017; Somerville et al., 2020, 2019). Given its high cost and nuanced benefits, biochar should be used sparingly to improve urban soils.

## Supporting information

Supplemental File for Wagner et al. Top dressed biochar increases tree seedling growth and decreases sodium leaching

## ACKNOWLEDGEMENTS

We thank the many people who assisted in experiment establishment and deconstruction and sample analyses: Michelle Catania and volunteers in the Soil Ecology Lab at The Morton Arboretum helped establish the project, conduct destructive sampling, and collect growth and biomass measurements; and Sav Henderson and Abby Malatia conducted a subset of the sodium leachate analyses. We also thank Plant Production staff at The Morton Arboretum who provided and maintained the greenhouse facility.

