## Supplemental File for Wagner et al. Top dressed biochar increases tree seedling growth and decreases sodium leaching for "Top dressed biochar increases tree seedling growth and decreases sodium leaching"

Supplemental Information

**Supplemental Table 1.** F and p values for SPAD, photosynthesis rate, and various growth responses of *Acer saccharum* to soil treatment (control, top dressed biochar, and incorporated biochar), salt treatment (no salt added and salt added), and interaction of soil and salt treatments.

| **Response** | **Soil treatments** | **Salt addition** | **Soil x salt** |
| --- | --- | --- | --- |
| SPAD | F_2,28.56_ = 1.77,  *p* = 0.19 | F_1,28.60_ = 0.03,  *p* = 0.86 | F_2,28.56_ = 3.10,  *p* = 0.06 |
| Photosynthesis rate  (µmol C m^-2^ ^s-1^) | F_2,26.17_ = 1.24,  *p* = 0.30 | F_1,26.71_ = 0.43,  *p* = 0.52 | F_2,26.17_ = 3.82,  *p* = 0.04 |
| Diameter growth (%) | F_2,30_ = 1.43,  *p* = 0.25 | F_1,30_ = 0.55,  *p* = 0.47 | F_2,30_ = 1.66,  *p* = 0.21 |
| Stem growth (%) | F_2,36_ = 0.46,  *p* = 0.63 | F_1,36_ = 0.21,  *p* = 0.65 | F_2,36_ = 2.88,  *p* = 0.07 |
| Coarse root growth (%) | F_2,36_ = 1.06,  *p* = 0.36 | F_1,36_ = 0.63,  *p* = 0.43 | F_2,36_ = 1.32,  *p* = 0.28 |
| New shoot biomass (g) | F_2,30_ = 0.71,  *p* = 0.50 | F_1,30_ = 1.24,  *p* = 0.27 | F_2,30_ = 0.96,  *p* = 0.39 |
| New fine root biomass (g) | F_2,36_ = 1.93,  *p* = 0.16 | F_1,36_ = 7.98,  *p* = 0.01 | F_2,36_ = 0.43,  *p* = 0.65 |
| Stem and coarse root biomass (%) | F_2,36_ = 0.80,  *p* = 0.46 | F_1,36_ = 0.06,  *p* = 0.81 | F_2,36_ = 0.13,  *p* = 0.88 |
| New shoot and fine root biomass (g) | F_2,30_ = 1.37,  *p* = 0.27 | F_1,30_ = 4.25,  *p* = 0.05 | F_2,30_ = 0.94,  *p* = 0.40 |
| Total biomass (%) | F_2,36_ = 0.90,  *p* = 0.42 | F_1,36_ = 0.30,  *p* = 0.59 | F_2,36_ = 0.16,  *p* = 0.85 |

**Supplemental Table 2.** F and p values for SPAD, photosynthesis rate, and various growth responses of *Gleditsia triacanthos* to soil treatment (control, top dressed biochar, and incorporated biochar), salt treatment (no salt added and salt added), and interaction of soil and salt treatments.

| **Response** | **Soil treatments** | **Salt addition** | **Soil x salt** |
| --- | --- | --- | --- |
| SPAD | F_2,9.48_ = 0.04,  *p* = 0.97 | F_1,9.85_ = 1.98,  *p* = 0.19 | F_2,911.87_ = 0.79,  *p* = 0.48 |
| Photosynthesis rate  (µmol C m^-2^ ^s-1^) | F_2,2.09_ = 3.90,  *p* = 0.20 | F_1,2.11_ = 3.77,  *p* = 0.18 | F_2,2.34_ = 3.62,  *p* = 0.19 |
| Diameter growth (%) | F_2,36_ = 2.76,  *p* = 0.08 | F_1,36_ = 0.05,  *p* = 0.82 | F_2,36_ = 1.33,  *p* = 0.28 |
| Coarse root growth (%) | F_2,30_ = 3.33,  *p* = 0.05 | F_1,30_ = 0.01,  *p* = 0.91 | F_2,30_ = 0.50,  *p* = 0.61 |
| New shoot biomass (g) | F_2,36_ = 1.98,  *p* = 0.15 | F_1,36_ = 0.71,  *p* = 0.41 | F_2,36_ = 0.22,  *p* = 0.80 |
| New fine root biomass (g) | F_2,36_ = 2.50,  *p* = 0.10 | F_1,36_ = 1.44,  *p* = 0.24 | F_2,36_ = 0.49,  *p* = 0.62 |
| Stem and coarse root biomass (%) | F_2,30_ = 1.30,  *p* = 0.29 | F_1,30_ = 0.24,  *p* = 0.63 | F_2,30_ = 0.49,  *p* = 0.62 |
| New shoot and fine root biomass (g) | F_2,36_ = 2.78,  *p* = 0.12 | F_1,36_ = 1.06,  *p* = 0.31 | F_2,36_ = 0.24,  *p* = 0.79 |
| Total biomass (%) | F_2,30_ = 3.62,  *p* = 0.04 | F_1,30_ = 0.01,  *p* = 0.92 | F_2,30_ = 0.59,  *p* = 0.56 |

**Supplemental Table 3.** F and p values for SPAD, photosynthesis rate, and various growth responses of *Quercus rubra* to soil treatment (control, top dressed biochar, and incorporated biochar), salt treatment (no salt added and salt added), and interaction of soil and salt treatments.

| **Response** | **Soil treatments** | **Salt addition** | **Soil x salt** |
| --- | --- | --- | --- |
| SPAD | F_2,23.15_ = 2.20,  *p* = 0.13 | F_1,22.39_ = 0.05,  *p* = 0.83 | F_2,21.74_ = 3.09,  *p* = 0.07 |
| Photosynthesis rate  (µmol C m^-2^ ^s-1^) | F_2,23.65_ = 1.61,  *p* = 0.22 | F_1,22.46_ = 0.01,  *p* = 0.92 | F_2,21.70_ = 2.98,  *p* = 0.07 |
| Diameter growth (%) | F_2,35_ = 0.72,  *p* = 0.49 | F_1,35_ = 2.20,  *p* = 0.15 | F_2,35_ = 0.05,  *p* = 0.95 |
| Stem growth (%) | F_2,35_ = 1.29,  *p* = 0.29 | F_1,35_ = 1.14,  *p* = 0.29 | F_2,35_ = 0.10,  *p* = 0.90 |
| Coarse root growth (%) | F_2,29.41_ = 0.06,  *p* = 0.94 | F_1,29.41_ = 2.56,  *p* = 0.12 | F_2,29.41_ = 1.03,  *p* = 0.37 |
| New shoot biomass (g) | F_2,36_ = 0.41,  *p* = 0.67 | F_1,36_ = 0.27,  *p* = 0.61 | F_2,36_ = 1.88,  *p* = 0.17 |
| New fine root biomass (g) | F_2,36_ = 0.62,  *p* = 0.54 | F_1,36_ = 0.06,  *p* = 0.82 | F_2,36_ = 0.84,  *p* = 0.44 |
| Stem and coarse root biomass (%) | F_2,29.05_ = 1.97,  *p* = 0.16 | F_1,29.05_ = 0.01,  *p* = 0.91 | F_2,29.05_ = 0.54,  *p* = 0.59 |
| New shoot and fine root biomass (g) | F_2,30_ = 0.74,  *p* = 0.49 | F_1,30_ = 0.13,  *p* = 0.72 | F_2,30_ = 1.41,  *p* = 0.26 |
| Total biomass (%) | F_2,35_ = 1.52,  *p* = 0.23 | F_1,35_ = 0.31,  *p* = 0.58 | F_2,35_ = 0.83,  *p* = 0.44 |

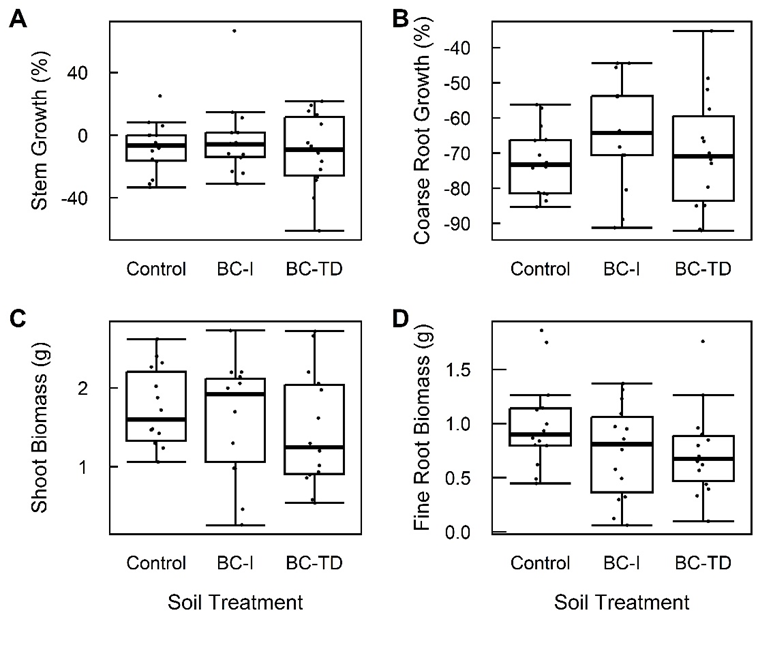

**Supplementary Figure 1.** Box and whisker plots of *Acer saccharum* percent stem growth (**A**), percent coarse root growth (**B**), shoot biomass (**C**), and fine root biomass (**D**) by soil treatment. The thick horizontal line inside each box represents the median value, the box represents the 25-75 percentile, and the whiskers extend from the first quartile minus 1.5*the interquartile range to the third quartile plus 1.5*the interquartile range. Each point on the graph is an individual tree’s sodium leachate measurement from that week. Control = control soils (no biochar), BC-I = incorporated biochar, BC-TD = top dressed biochar.

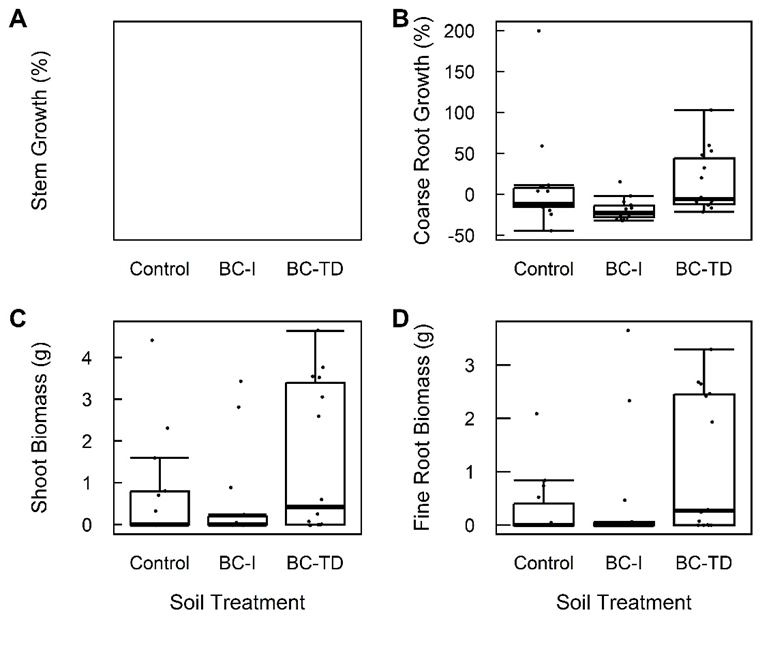

**Supplementary Figure 2.** Box and whisker plots of *Gleditsia triacanthos* percent stem growth (**A**, no data), percent coarse root growth (**B**), shoot biomass (**C**), and fine root biomass (**D**) by soil treatment. The thick horizontal line inside each box represents the median value, the box represents the 25-75 percentile, and the whiskers extend from the first quartile minus 1.5*the interquartile range to the third quartile plus 1.5*the interquartile range. Each point on the graph is an individual tree’s sodium leachate measurement from that week. Control = control soils (no biochar), BC-I = incorporated biochar, BC-TD = top dressed biochar.

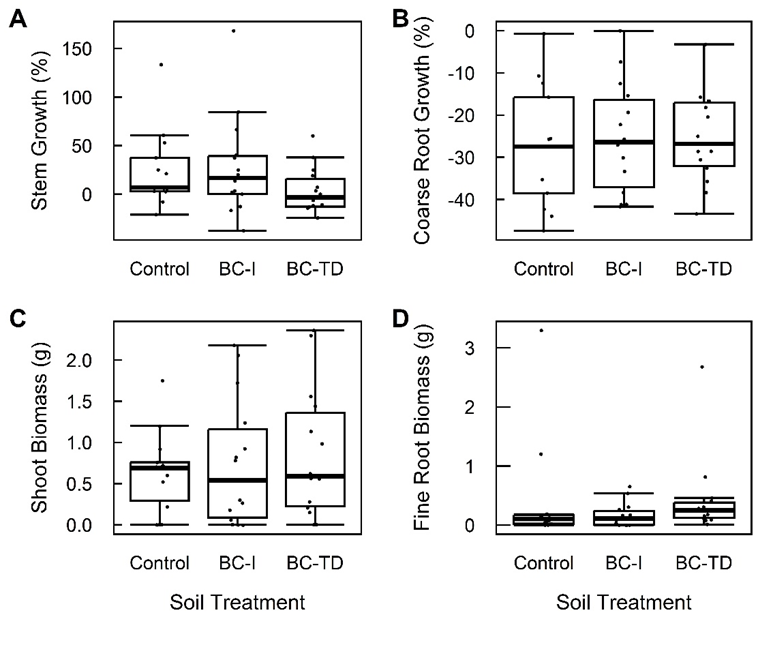

**Supplementary Figure 3.** Box and whisker plots of *Quercus rubra* percent stem growth (**A**), percent coarse root growth (**B**), shoot biomass (**C**), and fine root biomass (**D**) by soil treatment. The thick horizontal line inside each box represents the median value, the box represents the 25-75 percentile, and the whiskers extend from the first quartile minus 1.5*the interquartile range to the third quartile plus 1.5*the interquartile range. Each point on the graph is an individual tree’s sodium leachate measurement from that week. Control = control soils (no biochar), BC-I = incorporated biochar, BC-TD = top dressed biochar.
